# Topographic preservation of functional connectivity among the association areas in the human cerebral cortex

**DOI:** 10.1101/639252

**Authors:** Naoko Koide-Majima, Shinji Nishimoto

## Abstract

In the human sensorimotor cortex, some long-range corticocortical connections appear to preserve a fine-scale topology, in which physically close locations in the cortical region are functionally connected to physically close locations in other cortical regions. However, little is known about whether such topography preservation is unique to the sensorimotor areas or is general across other cortical areas. To investigate this question, we measured voxel-level functional connectivity using functional magnetic resonance imaging (fMRI) and visualized the fine-scale spatial organization of the connectivity patterns across the cortical surface. We found topographical preservation across regions, including the default mode network. Our results suggest that the topographical preservation of functional connectivity is not restricted to the sensorimotor cortex but also occurs in the association cortex.

## Introduction

Brain representations tend to change smoothly across the human cerebral cortex [Huth et al., 2012, Marglies et al., 2016]. In the early visual cortex, regions with similar representations, such as selectivity for similar visual angle or eccentricity, are located close to each other [Engel et al., 1997, Hansen et al., 2007]. Similar smooth gradients have been observed in other areas, including tonotopic representations in the auditory cortex [Da Costa et al., 2011, Langers and van Dijk, 2011, Leaver and Rauschecker 2016], somatotopic representations in the primary somatosensory cortex [Schieber, 1999, Schubotz and von Cramon, 2003], and category representation in the association areas [Huth et al., 2012]. Huth et al., 2016 [Huth et al., 2016] reported that the representation of speech-content changes smoothly from sensory to more abstract representation across the temporoparietal junction (TPJ), which is an association area within the default mode network (DMN) [Greicius et al., 2003, Fox et al., 2005].

Some of these smooth representational gradients have been linked to the topographic organization of connectivity patterns between distinct brain regions. These patterns are known as connectopic patterns, in which neighboring locations in a cortical region are connected with neighboring locations in a distant region [Thivierge and Marcus, 2007]. This type of organization has been observed in the human sensorimotor cortex: blood oxygen level-dependent (BOLD) signals of V1 voxels shown to predict signals in V3 voxels at similar locations in the retinotopic map than those at dissimilar locations [Heinzle et al., 2011]. In the auditory [Cha et al., 2014] and somatosensory cortices [Zeharia et al., 2015, Cauda et al., 2011], corresponding regions in the right and left hemispheres with similar representations (tonotopic and somatotopic representation) are functionally connected. However, such topographical preservation of connectivity has not been widely observed in the association areas, even though smooth gradients of representation have previously been observed in one association area—the TPJ [Huth, 2016].

Functional connectivity patterns have been often investigated by measuring the correlation of activity of brain regions during a resting-state task [Heinzle et al., 2011, Cha et al., 2014, Zeharia et al., 2015, Cauda et al., 2011, Koch et al., 2002, Hagmann et al., 2008, Honey et al., 2009, Tao et al., 2015]. Recent studies have demonstrated that tasks such as viewing movies produce more reliable identification of functional connectivity patterns than resting-state conditions [Vanderwal et al., 2017, Finn et al., 2017, Wang et al., 2017]. This reliability is independent of the resolution of cortical parcellation or the duration of the data collection used to calculate functional connectivity [Vanderwal et al., 2017]. Less head motion was observed in a movie-viewing task than in a resting-state task [Vanderwal et al., 2015], possibly leading to higher-quality-data acquisition in a movie-viewing task. Therefore, a naturalistic viewing paradigm considered to be an efficient approach for the measurement of fine-scale functional connectopic patterns.

In this study, we addressed the question whether a topographic organization of functional connectivity (fcTO) is present in the association areas. We specifically hypothesized that fcTO exists between the TPJ and the association areas in the DMN. We used functional magnetic resonance imaging (fMRI) to measure BOLD signals while eight subjects watched movie clips. To quantify the intensity of connectivity at a fine scale, we calculated the correlation between BOLD signals in each voxel in the TPJ, and each voxel in the external regions. To visualize the organization of the connectivity, we labeled each voxel in the external regions using the spatial coordinates of the strongly-connected voxels in the TPJ (see Supplemental Information). By visualizing the cortical map of the spatial coordinates, we demonstrated fcTO preservation within the DMN. We also performed a comprehensive automated search for fcTO for 156 anatomically-defined target regions across the cortex. We detected fcTO across several networks, including the DMN-related areas.

## Results

To capture the topographic organization of functional connectivity (fcTO), we developed a method for visualization of the structure of fine-scale connectivity structure between a seed region and external regions in the cerebral cortex (Figure 1). We focused on one seed region (e.g., the right transverse occipital sulcus), and labeled each voxel in the seed region with its spatial coordinate. For each voxel in the external region we calculated the strength of the connection to each voxel in the seed region using Pearson’s correlation coefficient between the BOLD-signal time courses [Kahnt et al., 2012, Cauda et al., 2011]. The voxel in the external region was labeled with the representative spatial coordinates of strongly-connected seed voxels (see Supplemental Information). If an external region showed a spatial gradient of labels with a similar topological order of the seed spatial coordinates, fcTO is considered to be preserved between the external region and the seed region. To visualize the spatial coordinates of the seed region and the external region, we assigned different colors to the spatial coordinates on the cortical connectopic map. An example of a cortical map was shown in Figure 1B. In this map connections are indicated by color: voxels in the seed region with a specific color (e.g., turquoise) had been shown to have a strong connection with voxels in the external regions represented by the same color.

**Figure 1.**
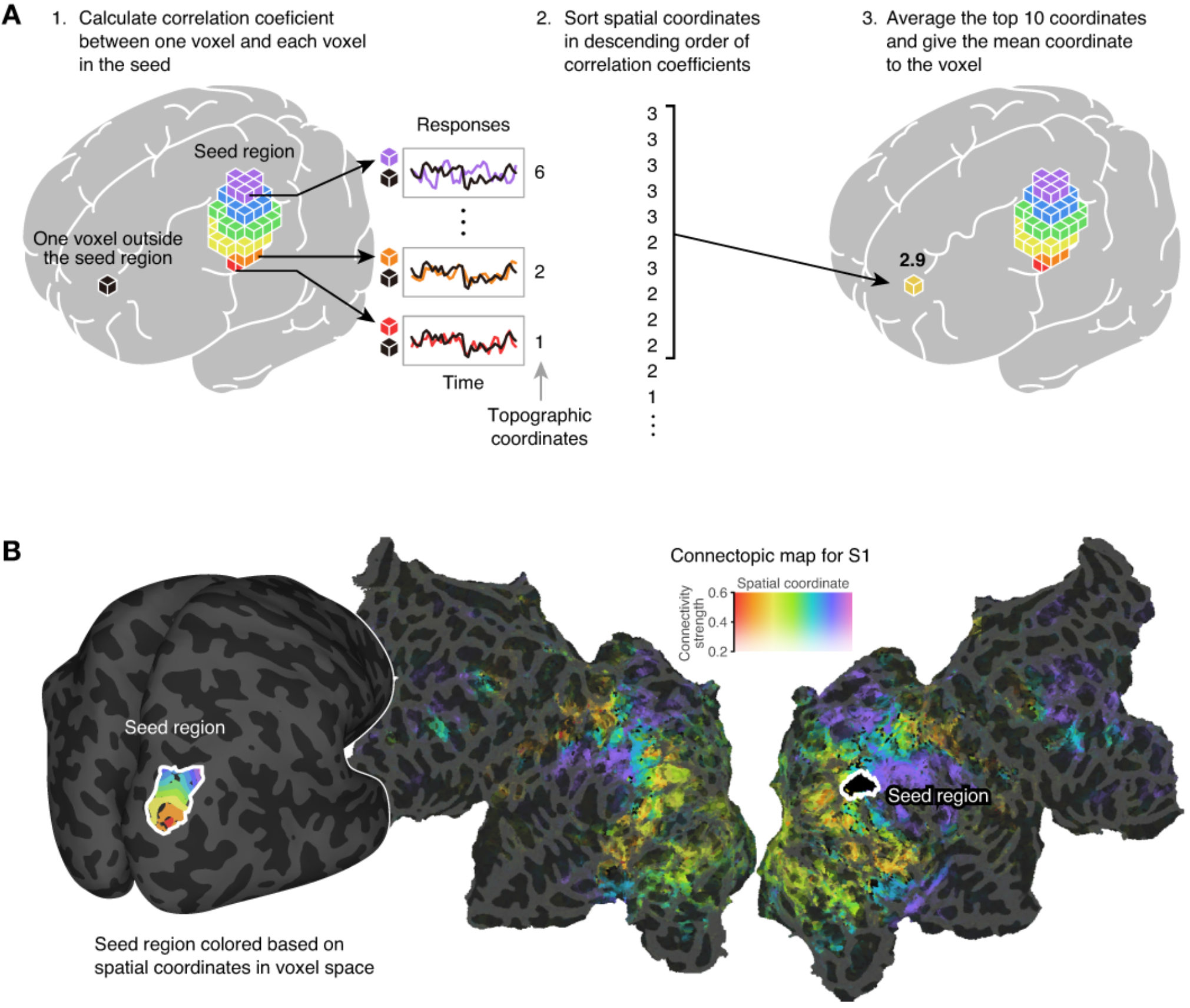
Workflow for visualization of a fine-scale connectopic map, given a seed region. (A) Topography in the seed region is defined using spatial coordinates and the coordinate assigned to a connected voxel in the external region. The connectopic map is a cortical map consisting of the spatial coordinates. (B) Example of a connectopic map when the transverse occipital area was used as a seed region. (Left) The location of the seed region on an inflated cortex. (Right) A connectopic map on the flattened cortical sheets for subject S1. Colored regions in the external (non-seed) areas show strong functional connectivity with the locations in the seed region of the corresponding color (Left). The opacity of the connectopic map represents connectivity strength (correlation) with the seed region.

### Topographic organization was preserved between the manually-defined TPJ and distant external regions

To identify fcTO preservation in the association areas, we focused on one region: temporoparietal junctions (TPJ). The TPJ is one of the transmodal regions [Marglies et al., 2016] and is a well-known region within the DMN [Fox et al., 2005]. We manually defined the left TPJ for each subject, and visualized its connectopic map (Figure 2, see Supplemental Information). The connectopic map of one subject, S1, showed strong functional connectivity between TPJ and several distant regions within the bilateral DMN (Figure 2A). Some regions exhibited color gradients that preserve the color orders of the seed region, including the right TPJ, the bilateral middle temporal cortex, the precuneus, and the frontal cortex. This observation suggests that there is a clear fcTO between these regions. We observed similar topographic patterns in the connectopic map for all of the other subjects (S2–S8) (Figure 2B, Figure S1).

**Figure 2.**
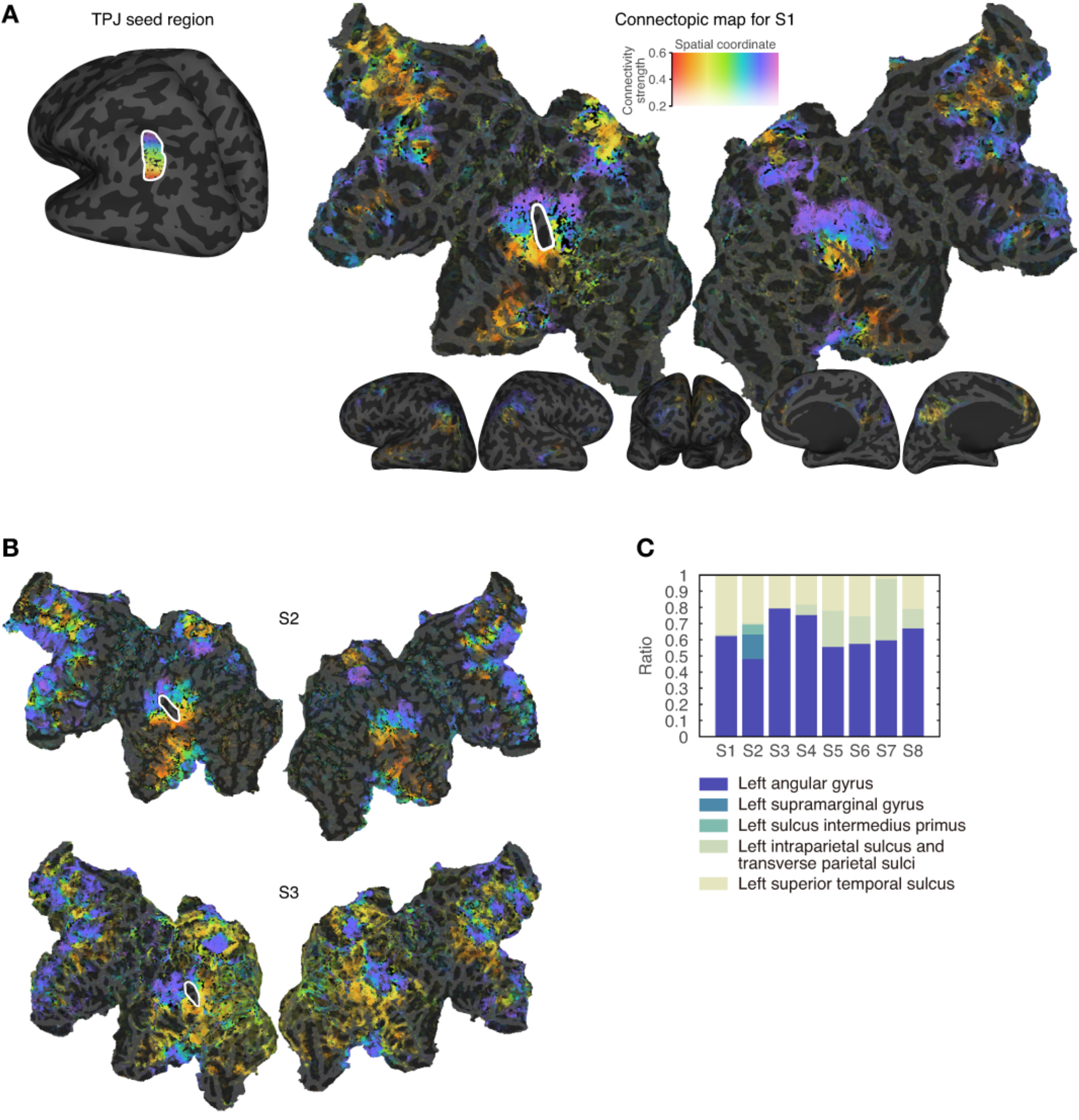
Connectopic map of the TPJ showing topography preservation in functional connectivity to the external regions. (A) (left) The manually-defined TPJ seed region. (right) The connectopic map shows several distinct regions that show strong connections to the TPJ. Several regions showed spatial color gradients that preserve the color order of the seed region, suggesting topographical preservation of the functional connectivity. (B) Connectopic maps for two other two subjects (S2 and S3; see Figure S1 for all other subjects). In these maps, the gradients are similar to those of S1. (C) Voxel ratio of anatomical labels of the Destrieux atlas [Destrieux et al., 2010] in the TPJ for each subject. Each seed consists mainly of the angular, the Intraparietal and the superior temporal sulcus, which is consistent across all subjects.

### Quantitative verification of topographic functional connectivity

To quantitatively examine the fcTO preservation between the TPJ and external regions, as well as its reproducibility across subjects, we manually defined seven representative lines of interest in the external regions where: (1) the connectivity with TPJ is high; (2) visible spatial gradients are observed; and (3) both (1) and (2) are preserved across subjects (Figures 1, 3A, and Figure S1). We extracted spatial coordinates along each line and compared these coordinates across the subjects (solid lines in Figure 3B). To examine whether the spatial coordinates on the line have a similar topological order as those observed in the TPJ (diagonal gray-dashed line in Figure 3B), we calculated the mean squared error (MSE) between the normalized spatial coordinates on the line and those for the TPJ (see Supplemental Information). The MSE was significantly lower than the variance of the spatial coordinates on the line (line 1: *F*_154_ = 2.07; line 2: *F*_159_ = 2.17; line 3: *F*_159_ = 3.10; line 4: *F*_159_ = 2.57; line 5: *F*_159_ = 2.20; line 6: *F*_158_ = 2.70; line 7: *F*_159_ = 4.07; *p* < 0.01 after Bonferroni correction for all lines, using an F test). This indicates the preservation of the fcTO between the TPJ and the external regions.

**Figure 3.**
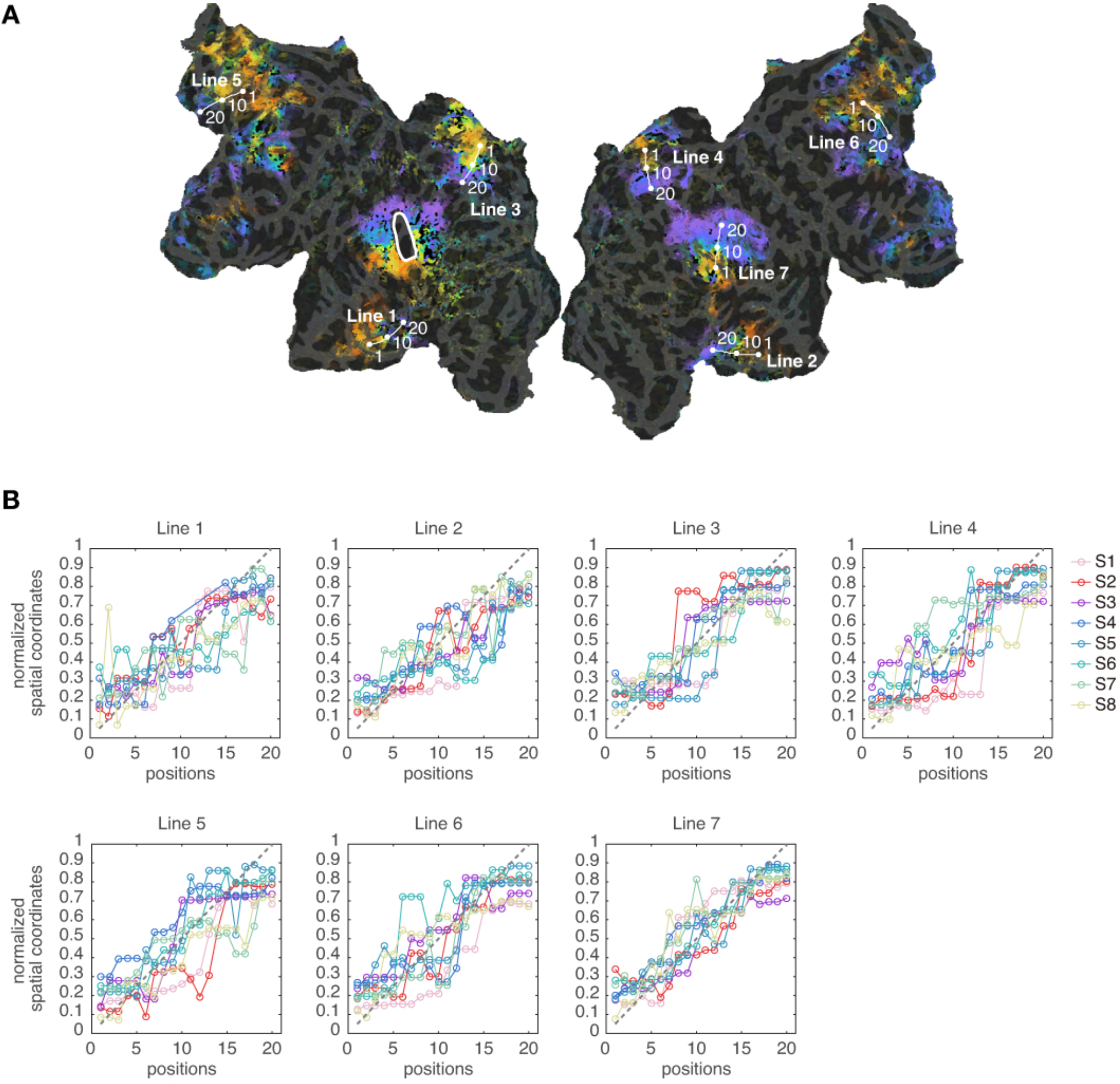
Region-wise examination of topography preservation across the subjects. (A) Visualization of a connectopic map of the TPJ with the seven representative lines of interest (S1). The lines were manually defined in regions showing high functional connectivity and a topography-preserving gradient of spatial coordinates with the TPJ (for the other subjects S2–S8 see Figure S2). (B) Spatial coordinates along the lines for all eight subjects. For all lines shown, spatial coordinates tend to monotonically increase (*p* < 0.01 with Bonferroni correction, F test), which supports that spatial coordinates on the line had a similar topological order of the TPJ coordinates (gray-dashed line).

### Effect of stimulus-induced BOLD signals on functional connectivity measurements

The connectopic map for the TPJ was created using BOLD signals measured during a natural-movie viewing task. It is possible that stimulus-induced co-activation of BOLD signals across voxels might be interpreted as functional connectivity. To investigate this possibility, we quantified the ratio of stimulus-induced signals (stimulus explainable variance, [Mante et al., 2008, Ikezoe et al., 2018]) using the reproducibility of the BOLD signals across four repetitions of identical movie stimuli (Figure S3). We found that the TPJ exhibited low-level stimulus-induced reproducibility. The average ratio of stimulus explainable variance was: S1:2.7%; S2:7.4%; S3:8.1%; S4:12.4%; S5:9.5%; S6:10.3%; S7:8.7%; S8:5.0%. This observation suggests that BOLD signals in the TPJ were largely driven by internal, non-stimulus, factors, and thus the fcTO preservation we found in the TPJ was not solely explained by the co-activation pattern of stimulus-locked BOLD signals.

### Automatic detection of fcTO-preserving regions

To further examine whether the fcTO preservation was specific to DMN-related regions, we developed a novel method for comprehensive automatic screening of regions showing fcTO preservation. A total of 156 candidate seed regions were determined using Destrieux atlas where the anatomical regions were defined across the whole cortex and sub-cortex (see Supplemental Information). We computed the connectopic map for each of the 156 candidate seed regions and used the spatial coordinates to automatically quantify the degree of fcTO preservation between the seed region and connected external regions, a metric dubbed topographic similarity (TS) (see Supplemental Information and Figure S4). A high TS indicates that the seed region preserves the fcTO with external regions. Figure 4A shows representative examples of connectopic maps for seed regions with low, middle, and high-TS values among the 156 candidate seed regions. These examples show that the high-TS region show clear fcTO preservation, while the topographic gradient was not clear in the middle-TS seed region and was not found in the low-TS seed region (Figure 4A and Figure S5).

**Figure 4.**
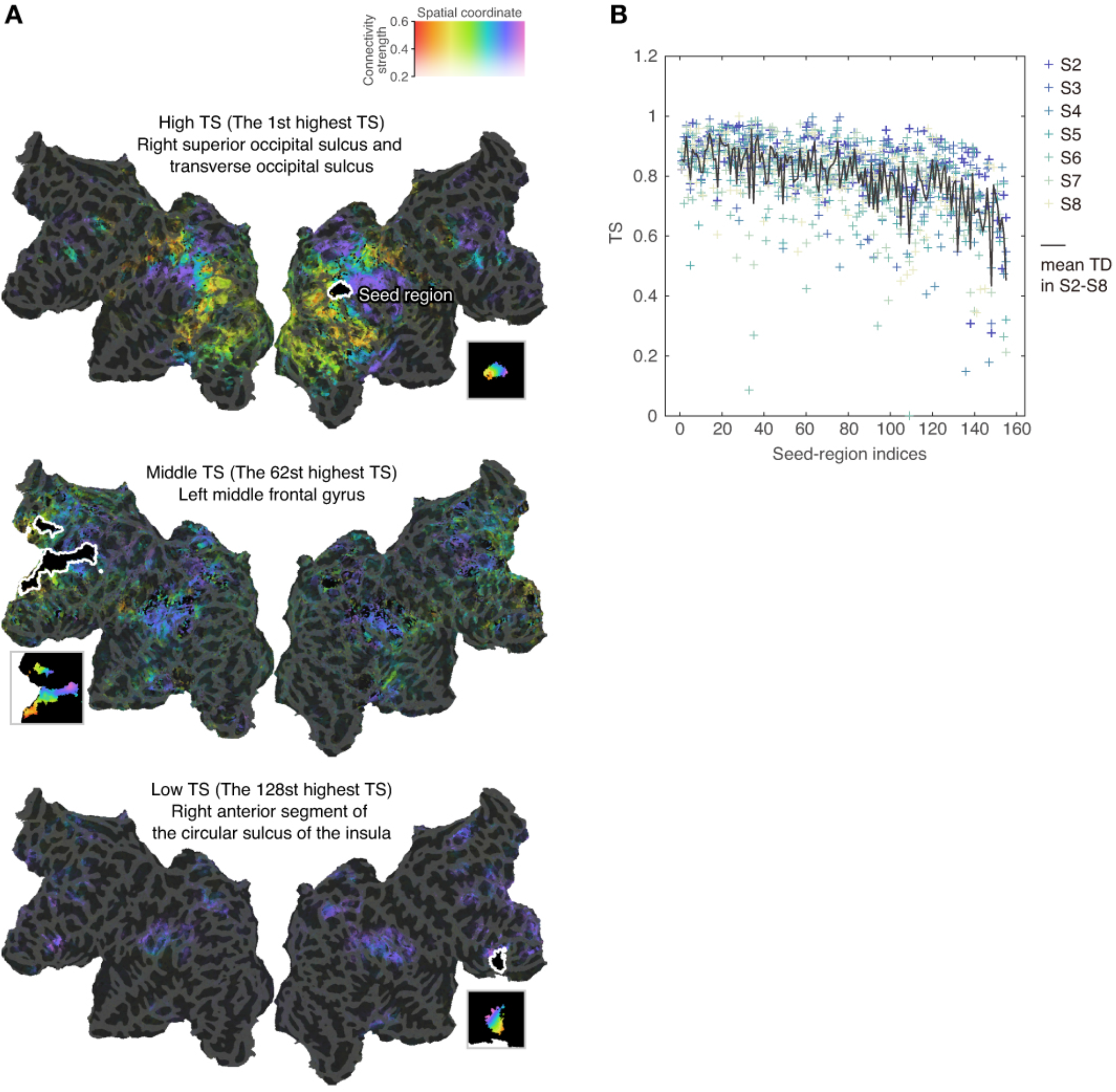
Topographic similarities (TSs) for the 156 anatomically-defined seed regions. High TS denotes the fcTO preservation between a seed region and the connected regions. (A) Examples of connectopic maps for a single subject, S1, for high, middle and low-TS seed regions across the 156 seed regions in the whole cortex and the sub-cortex. In the map of the high-TS seed region, we observe similar topographic gradients of the spatial coordinates between the seed region and the connected external regions. However, such gradient can not be observed in the middle and low-TS seed regions. (B) Plot of the TSs for the seven subjects (S2-S8) when seed regions are arranged in ascending order of TSs for a single subject (S1).

To confirm that the results of the analysis using TS values were consistent across subjects, we arranged the 156 seed regions in the ascending order of the TSs computed from a single subject (S1) and arranged the TSs computed from each of the seven remaining subjects (S2–S8) in the same order (Figure 4B). The TSs averaged across the seven subjects gradually decreased (*p* < 10^−47^, F test on the linear regression, Figure 4B).

To further examine which regions show high TS across subjects, we selected the seed regions with the top 30 TSs for each of the eight subjects. The seed regions selected in more than six subjects are listed in Table 1. We found that more than half of the selected seed regions are related to the TPJ, suggesting that the fcTO preservation was found more frequently in the TPJ than in the other regions.

**Table 1.**
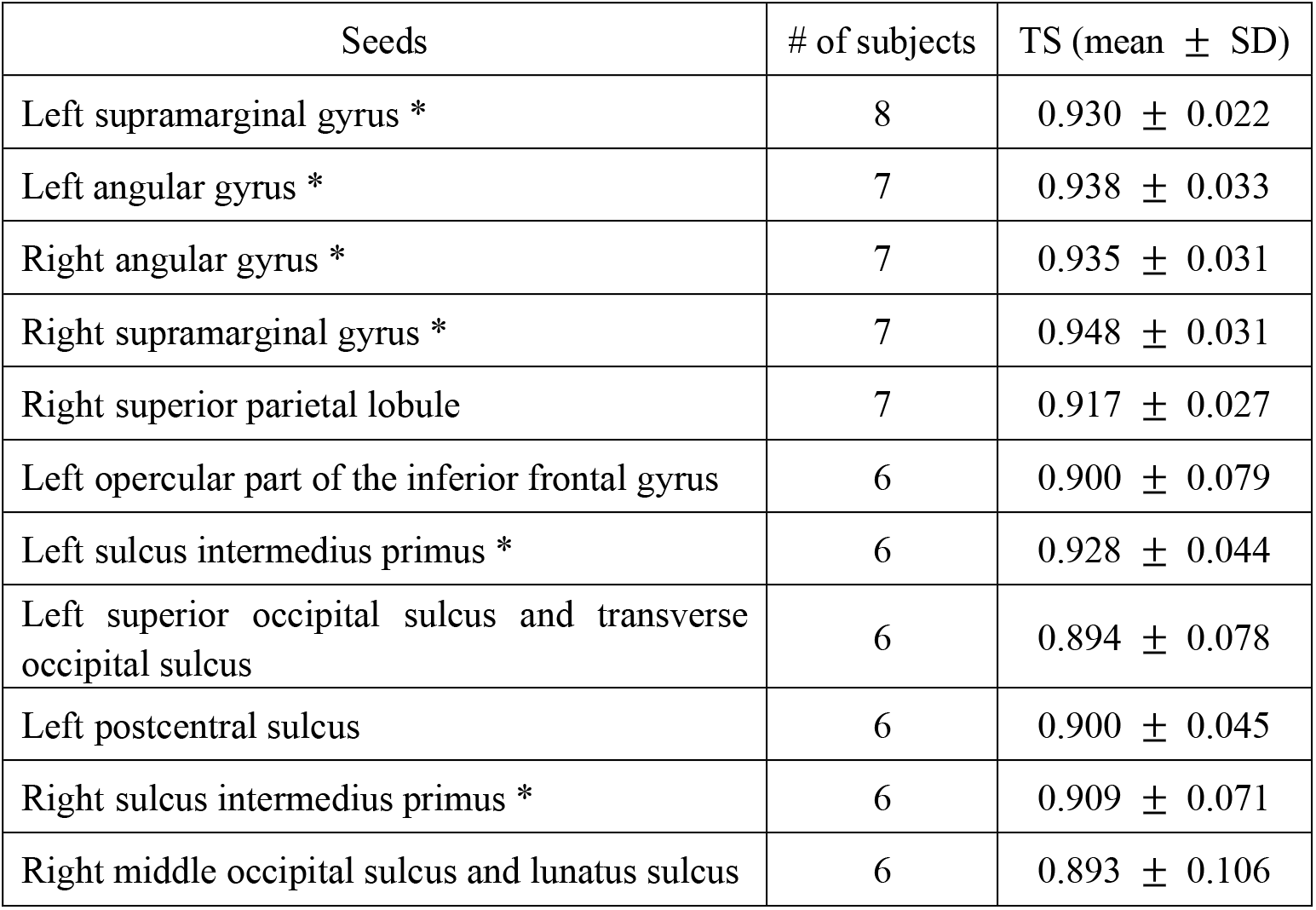
Seed regions with consistently high topographic similarities (TS) across subjects. TS was quantified as order similarity of spatial coordinates between the seed and those in the connected regions (see Supplemental Information). High TS denotes the fcTO preservation between a seed region and the connected regions. For each subject, we defined high-TS seed regions as those having the highest 30 TSs across the 156 seed regions in the whole cortex and sub-cortex. To find high-TS seed regions that were shared across subjects, we listed the seed regions that were consistently selected across more than six subjects (We had eight subjects). Seed regions marked with (*) denote regions within the temporoparietal junction.

## Discussion

Previous neuroimaging studies have demonstrated topography preservation between distant cortical regions; that is nearby locations in one region are functionally connected to nearby locations in another region [Heinzle et al., 2011, Cha et al., 2014, Zeharia et al., 2015, Cauda et al., 2011]. However, topography preservation was reported only for the primary sensorimotor cortex, and it is still unknown whether such topographic organization is ubiquitous throughout the cerebral cortex. We found that topographic organization in functional connectivity (fcTO) was preserved between the DMN-related regions (Figure 2, 3, Figure S1, S5), which are known to be related to transmodal representation [Margulies et al., 2016]. We also found fcTO preservation within the DMN, using our novel method for automatic detection of the fcTO-preserving regions through the whole cortex (Table 1, Figure 4, Figure S5, see Supplemental Information).

Topography preservation within the DMN may play an important role in efficient communication between the regions. It is assumed that evolutionary pressures act to minimize neural wiring [Buzsáki et al., 2004]. Previous diffusion imaging studies have reported a substantial correspondence between structural connectivity and resting-state DMN [Hagmann et al., 2008, Honey et al., 2009, Tao et al., 2015]. Collectively, these observations indicate the existence of straight wiring between the DMN-related regions, which leads to the fcTO preservation between the regions. Our results provide important data to assist with understanding the fundamental organization of the human cortex.

The fcTO preservation among the DMN-related regions might imply the existence of a functional map in the association areas, such as the retinotopic maps and the tonotopic maps in the primary sensory cortex [Engel et al., 1997, Hansen et al., 2007, Da Costa et al., 2011, Langers and van Dijk, 2011, Leaver and Rauschecker, 2016]. The DMN-related regions showed reduced brain activity during a task requiring attention to targets [Raichle et al., 2001, Buckner et al., 2008] and were considered to be related to an internally-focused task such as self-relevant memory retrieval and theory of mind [Buckner et al., 2008]. Recent studies have examined further details about the functions of the DMN-related regions [Spreng et al., 2010, Spreng, 2012, Huth et al., 2016]. In the DMN-related regions and their neighboring regions, smooth gradients from sensory-related to more abstract representation have been shown. For example there is a gradient from visuospatial planning to autobiolographical planning [Spreng et al., 2010, Spreng, 2012] and from visual and tactile concepts to those related to social and emotion during listening natural speech [Huth et al., 2016]. Our results regarding fcTO preservation among the DMN-related regions might reflect such functional representation gradients of abstraction levels.

The novel method we developed enables us to automatically detect seed regions that have fcTO preservation with the other regions (Figure S4D), while previous studies demonstrated such preservation using a pre-defined seed [Marquand et al., 2017, Kahnt et al., 2012, Cauda et al., 2011, Roy et al., 2009, Li et al., 2013]. The detection capability of our methods may depend on the manually-chosen parameter used to define the spatial coordinates. In this study, we used the dorso-ventral axis to define the spatial coordinates and found that the TPJ was one of seed regions showing fcTO preservation with the connected regions. fcTO preservation was observed using the pre-determined dorso-ventral axis, while an axis with a different direction may also be efficient for detection of such preservation. Further examination of the fcTO preservation is needed using different axes, to enable us to indentify unknown fcTO preservation within the other regions.

## Supplemental Information

### Subjects

Eight healthy individuals (S1–S8; age 23–32; four females) with normal or corrected-to-normal vision participated in our experiments. All subjects accepted that and gave written informed consent. The experimental protocol was approved by the ethics and safety committees of the National Institute of Information and Communications Technology.

### Experimental design and stimuli

Subjects watched audio-visual stimuli during fMRI data acquisition. The stimuli consisted of 138 movie clips from the video-sharing site *Vimeo* (https://vimeo.com/jp), which contains various genres, such as human drama, fantasy and daily life scenes, and action movies. Movies were clipped to 10–20 seconds in length and the stimulus sequence was produced by combining the selected clips in random order.

The visual stimuli were presented at the center of a projector screen with 23.3 × 13.2 degrees of visual angle, at 30 Hz. The audio stimuli were presented through MR-compatible earphones with an appropriate volume level for each subject. The subjects were instructed to watch the clips naturally as is the case with watching TV shows in daily life.

### MRI data acquisition

MRI data were collected on a 3T Siemens Trio TIM scanner (Siemens, Germany) using a standard Siemens 32-channel volume coil and a multiband gradient echo-planar imaging sequence [Moeller et al., 2010] with the following parameters; TR = 2000 ms, TE = 30 ms, flip angle = 62°; voxel size = 2 × 2 × 2 mm, matrix size = 96 × 96, 72 axial slices, FOV = 192 × 192 mm, multiband factor 3. Anatomical data were collected on the same 3T scanner using T1-weighted MPRAGE with the following parameters; TR = 2530 ms, TE = 3.26 ms, flip angle = 9°, voxel size = 1 × 1 × 1 mm, matrix size = 256 × 256, 256 axial slices, FOV = 256 × 256 mm.

For each subject, fMRI data were collected in three separate sessions over three or four days. In all sessions, 18 movie-watching runs were performed, with a resting period was followed after each run. Each run lasted 610 seconds. The data used in our main analysis were collected in 12 runs of the 18 runs and consisted of different movie clips (total 7200 seconds). The remaining six runs were performed for validation of the data acquisition. Across the six validation runs, three types of 300-second movie sequences, consisting of different movie clips, were repeated four times (total 3600 seconds). In each of the 18 runs, the fMRI data acquired in the first 10 seconds was not used in our analyses.

### fMRI data preprocessing

We used the Statistical Parameter Mapping toolbox (SPM12, http://www.fil.ion.ucl.ac.uk/spm/software/spm12/) to preprocess the EPI data. EPI data were subjected to slice-timing correction and aligned to the first image from the first scan for each subject. For each voxel, the response was normalized by subtracting the mean response across all time points. We then removed low-frequency responses by subtracting the convolution results of a median filter with a 120-second time window for every 2-second time point. To define anatomical regions for each subject, using FreeSurfer (http://surfer.nmr.mgh.harvard.edu/), the cerebral cortex was segmented into 156 regions as defined by the Destrieux atlas [Destrieux et al., 2010]. The segmentation results were registered from T1 space to EPI space using a FreeSurfer function, and each voxel was given one anatomical label.

### Definition of spatial coordinates in each seed region

To quantify the topographic organization in a seed region, we assigned a spatial coordinate to each voxel in the seed region. For a manually-defined seed, we assigned 2 mm-interval coordinates on the flattened cortical map along a manually-defined direction. For the automatically-defined seed from the anatomical 156 regions, we assigned coordinates along the dorso-ventral axis of the EPI space (Figure 1A). The range of the coordinates was different across subjects and seed types, ranging from 1 to 3–41, across the 156 seeds, because the sizes and shapes of the seed regions were different.

### Quantification of topographic organization in voxel-level functional connectivity between a seed region and external regions

To identify topographical preservation between a seed region and external regions, we labeled each voxel in the external region by the locations of voxels in the seed region that showed strong functional connectivity with it. Functional connectivity was calculated using Pearson’s correlation coefficient between the time course of the BOLD signals for each of the seed voxels and those for each of the external voxels [Kahnt et al., 2012; Cauda et al., 2011]. We labeled voxels in the seed region with the spatial coordinates along the pre-defined axis. For quantification of the fcTO with the manually-defined TPJ, the axis used was manually defined on the flattened cortical map. For automatic detection of fcTO-preserving seed regions, we used the dorso-ventral axis. After labeling voxels in the seed region, we labeled each of the voxels in the external region with the spatial coordinates of a subset of strongly-connected voxels in the seed region. To label each of the external voxels, we selected the top 10 voxels in the seed region with respect to connectivity strength, and the averaged spatial coordinates of the 10 voxels. The averaged coordinate value was treated as the representative coordinate for the external voxel. However, this procedure does not always produce representative coordinates. If the spatial coordinates of the top 10 seed voxels have a large variance (e.g., [1, 9, 1, 1, 1, 9, 9, 1, 1, 9] from 1 to 10), the average coordinate (e.g., ‘4.2’) would not be representative of those voxels. To avoid such a case, we eliminated the representative coordinates from our results if those voxels showed a distance between the mode and the average of the top 10 coordinates of more than 20% range of spatial coordinates in the seed region. When calculating of the mode, when there were multiple values occurring with equally frequency, we treated the smallest of those values as the mode. For visualization of the representative spatial coordinates on the connectopic map, the coordinates are represented by a single color using a linear interpolation of the RGB values of the top 10 coordinates in the seed. To highlight the strongly-connected external voxels, the opacity of the coordinate colors was determined according to the connectivity strength. The connectivity strength of each external voxel was quantified as the averaged correlation coefficient across the top 10 seed voxels. Before averaging, a Fisher z-transformation was applied to the individual correlation coefficient values. The results were averaged across the 10 voxels and the averaged correlation coefficient was inversely z-transformed. The resulting value was used as the quantification of the connectivity strength.

### Manual definition of the TPJ and quantification of the topography preservation with the external regions

To find topography preservation within the DMN, we employed the left TPJ as a seed region. The seed region was manually defined on the flattened cortical map for each subject. Manual TPJ definition was performed on the flattened cortical map (e.g., 722 mm × 1326 mm for subject S1) to capture the position of the left angular gyrus of the Destrieux atlas [Destrieux et al., 2010]. The number of the voxels in the TPJ ranged from 165 to 235 (mean = 197.38, SD = 26.45) across the eight subjects. The TPJ was defined to be in the anatomically similar position across subjects. To support that, we summarized the anatomical labels in the TPJ for each subject (Figure 2C). In the results the TPJ included the angular, the intra-parietal and the superior temporal areas, consistently across all subjects.

To confirm whether the fcTO was preserved between the TPJ and the external regions, we manually defined seven representative lines with strong connections, and examined whether the spatial coordinates in the seed were significantly similar to those on each line (Figure 3A). Lines were defined on the flattened cortical map. The lines were defined firstly for a single subject S1. For each of the seven remaining subjects (S2–S8), the lines corresponding anatomically to S1’s lines were defined. We summarized the anatomical labels obtained from the regions upon which each line was laid on (Figure S2). As a result, at least one or two labels for each line were shared across all subjects, and the remaining three to six labels were neighbors to the shared labels (the total number of anatomical labels for all lines was 37). The seven regions corresponding to the seven lines were mainly as follows: the left and the right middle-temporal cortex for lines 1 and 2; the left and the right precuneus for lines 3 and 4; the left and the right superior-to-middle frontal cortex for lines 5 and 6; the right angular and superior temporal cortices for line 7. All regions were involved in the DMN. For each line, we set 20 even-interval positions on the line, and set a 6-mm-radius circle area at each position. At each position, we used the mode of the spatial coordinates in the circle area as the representative spatial coordinate. The spatial coordinates were normalized by subtracting the maximum number of spatial coordinates of the TPJ. The TPJ coordinates were also normalized in the same manner, with 20 coordinates at even intervals, linearly increasing from 0 to 1. To quantify the fcTO preservation between the seed region and each line, we calculated the mean squared error (MSE) between the normalized spatial coordinates on each line (Figure 3B, solid line plot) and the TPJ coordinates (Figure 3B, gray-dashed line plot) and tested whether the MSE was significantly lower than the variance of the spatial coordinates on the line.

### Automatic detection of seed regions showing topography preservation

To further examine potential fcTOs of arbitrary regions across the cortex, we developed an automated method to detect and quantify topographic similarity as described below (Figure S4).Briefly, each candidate seed region was taken from the 156 regions identified in the Destrieux atlas [Destrieux et al., 2010], which were anatomically defined using FreeSurfer (http://surfer.nmr.mgh.harvard.edu/). For each seed region, functionally connected areas were screened and quantified in terms of the degree of topographical preservations (Figure 4A, Figure S5).

We selected 15 fcTO candidate regions that were strongly connected to each seed region (Figure S4A). The 15 candidate regions were obtained as a circular area with a radius of 50 mm, roughly half the average area of an anatomically-defined seed region, large enough to capture the local spatial coordinates. The 15 candidate regions were obtained at 15 peaks of the connectivity strength (= correlation coefficients) on the flattened cortical sheet as follows. First, the cortical map of the connectivity strength was spatially smoothed using a Gaussian filter with a width of 4 mm to avoid detecting noisy local peaks. Then, the 15 peak locations were detected using an iterative process. The location with the highest connectivity was detected on the smoothed connectivity map as the first peak location (Figure S4A-1). Then, to avoid detecting overwrapping regions multiple times, we masked out a circular area with a radius of 140 mm at the first peak location (Figure S4A-2). Subsequently, we searched for the location with the highest connectivity in the remaining region as the second peak location. We again masked out a circle with a radius of 140 mm at the second peak, before searching for the third peak location. This detection step was repeated 15 times, resulting in 15 peak locations were identified as the candidate regions (Figure S4A-3).

To quantify the degree of topography preservation for each of the candidate regions, we set a representative curve line within each candidate region (Figure S4B). The curve line was optimized so that the spatial coordinates along the line would be as topographically linearly ordered as possible (Figure S4C) and were obtained using the following three steps. First we obtained the candidate endpoints of a representative curve line. The candidate endpoints were pairs of the three minimum-peak locations and the three maximum-peak locations of the connectivity-strength map within the region. The candidate peak locations were detected iteratively using the detect-and-mask procedure described above to find the global-peak locations, with a circular area of radius 24 mm was masked out at each peak location. At the second step, for each of the endpoint pairs, we obtained the optimal curve line via one intermediate location showing linearly-ordered spatial coordinates. We searched for such an intermediate location in the circular area at the midpoint between the endpoints (the circle radius was one third of the edge length). For each candidate intermediate location, we fitted a cubic spline curve and obtained the modes of the spatial coordinates from the 4-mm-radius circles at even-interval positions on the spline curve. We used only the spatial coordinates of voxels showing high connectivity strength, with the representative correlation coefficients greater than 0.2. We then calculated the difference between the obtained spatial coordinates (modes) and a linearly-ordered coordinates using MSE (as in Figure 3B). The MSE was calculated for each candidate intermediate location, and we used the intermediate location showing the lowest MSE to fit the optimal curve line for the endpoint pair. At the third step, we compared MSEs across the optimal curve lines for endpoint candidates, and employed the curve line showing the lowest MSE as a representative curve line for the candidate region. To verify whether the representative curve line showed low MSE, we compared the MSE with those for randomly-defined 300 curve lines. As a result, the representative curve line had significantly lower MSE than those for 300 randomly-defined lines (*p* < 0.005, 300 permutations, Figure S4D).

We identified the five regions with the lowest MSEs across the 15 candidate regions as the five external regions that had high-order similarity to the seed region. To compute the TS, the MSEs of the five external regions were averaged, and the averaged MSE was normalized by dividing it by a variance of the spatial coordinates for the seed. Then the normalized MSE was inverted and re-normalized from 0 to 1 across all subjects and all seed regions. The re-normalized result was used as the TS for the seed region.

### Consistency in topography-preserving seed regions across subjects

For the automatic-detection of the fcTO-preserving seed regions, we calculated the TSs for the 156 anatomically-defined regions. To confirm that the fluctuation in TSs across the 156 seed regions was consistent across all subjects, we examined whether the high-TS seeds for a single subject also showed high TSs for the seven remaining subjects. For each of the seven subjects (S2-S8), we arranged 156 seed regions in descending order of the TSs in the first subject (S1, Figure 4B) and confirmed that TSs decreased as the arranged-region order increased. To find seed regions that had high TSs consistently across subjects, for each subject, we selected the seed regions having the 30 highest TSs across the 156 seed regions. We then counted the number of subjects in which a given region was selected. The number ranged from one to eight, with the number eight denoting that the seed region was shown in the top 30 seed regions across all subjects. We listed seed regions with more than six occurences in Table 1.

## Acknowledgments

This study was partly funded by Brother Industries Ltd. and JSPS KAKENHI Grant Numbers 15H05311 and 15H05710 as well as JST CREST JPMJCR18A5 and ERATO JPMJER1801. Data were collected with support from the National Institute of Information and Communications Technology.

## Author contributions

N.K. and S.N. designed the experiment; N.K collected the data; N.K analyzed the data with support from S.N.; N.K. and S.N. wrote the manuscript.

## Data availability

The data that support the findings of this study are available from the corresponding author upon request.

---

**Figure S1.**
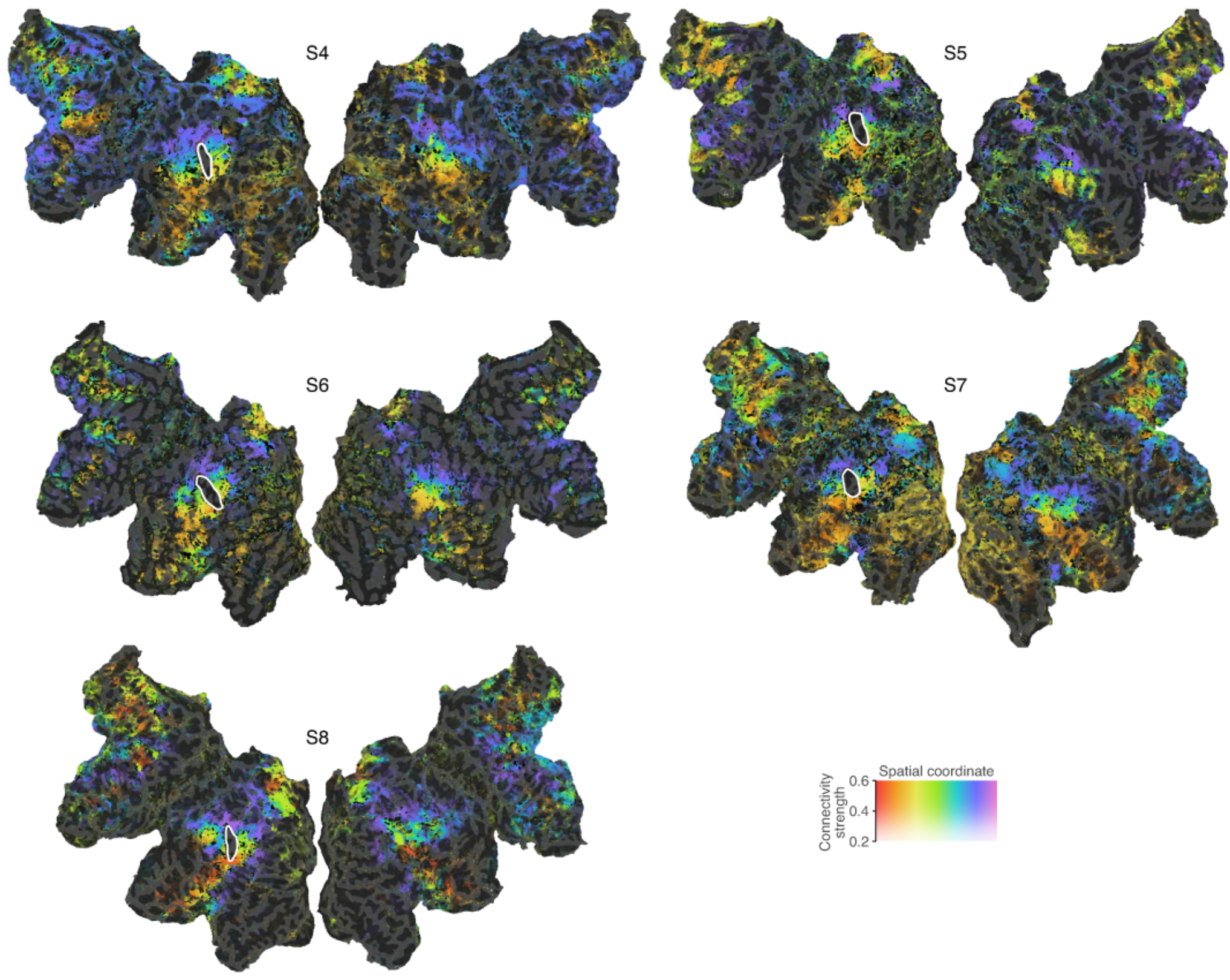
Connectopic maps of the manually-defined TPJ for the five subjects (S4–S8). For all maps, similar topographic gradients were observed in the TPJ and the connected external regions, consistently across the five subjects and the other three subjects in Figure 3. Related to Figure 2.

**Figure S2.**
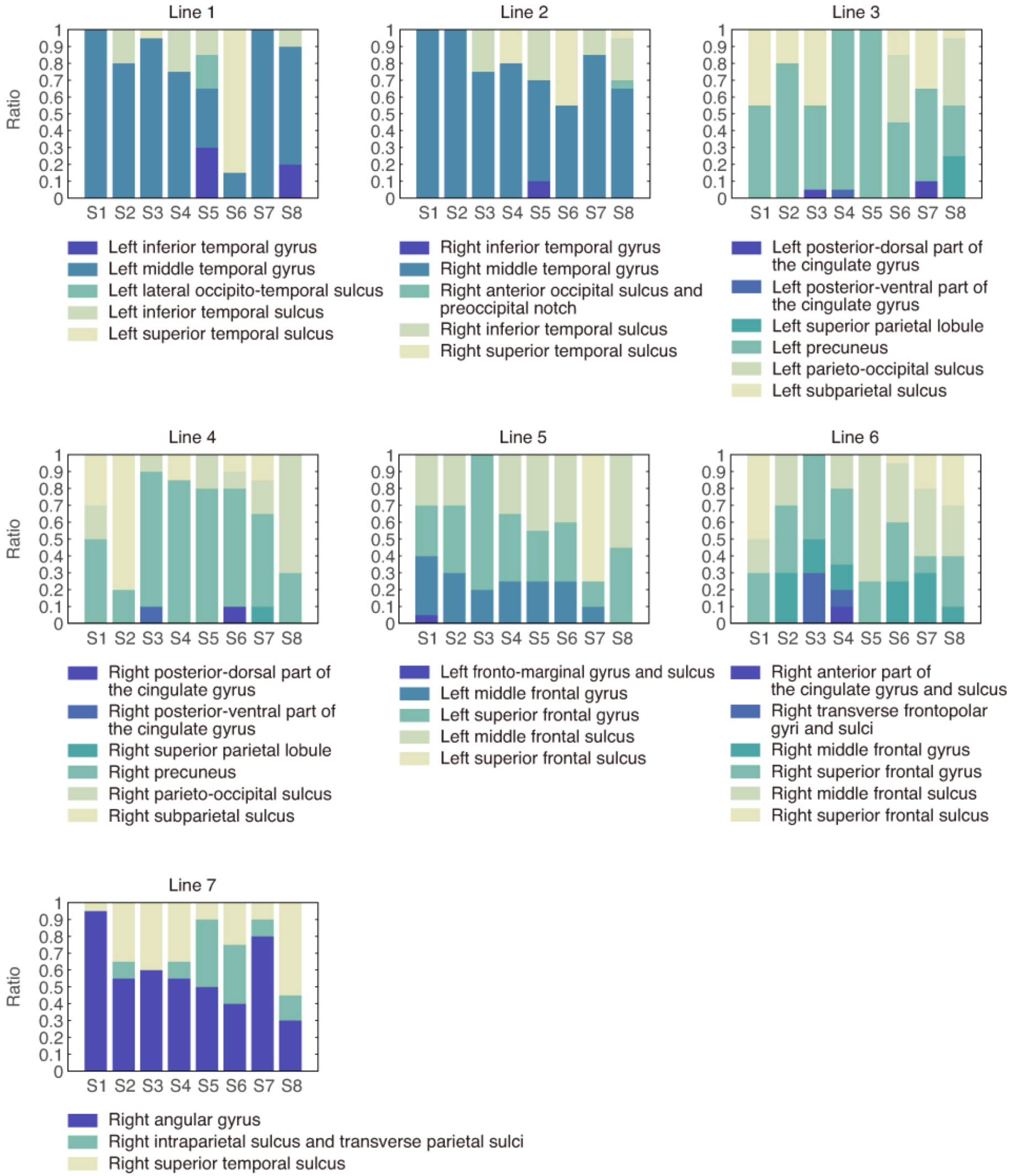
Area ratios of anatomically-defined labels for the seven regions connected to the manually-defined TPJ. For each subject, we obtained 20 labels from 20 positions on the line in each of the connected regions and counted the frequency for each of the unique labels across the 20 labels. For each line, the same or neighboring anatomical labels were included across all subjects. The anatomical labels for all lines were known to be regions within the default mode network [Fox et al., 2005]. Related to Figure 3.

**Figure S3.**
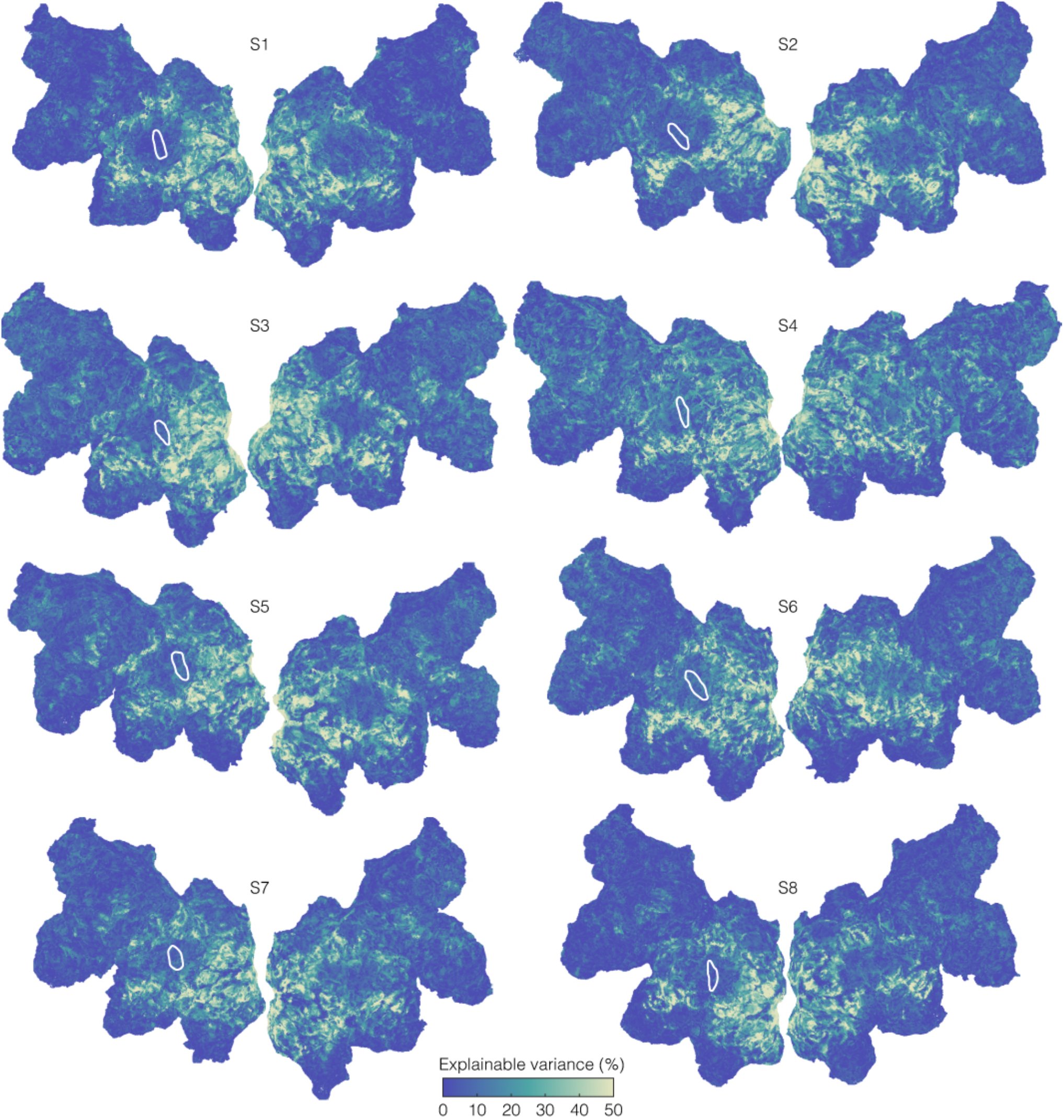
Reproducibility of BOLD signals across four repetitions of movie stimuli. Reproducibility was quantified as an explained variance (EV) [Mante et al., 2008, Ikezoe et al., 2018], and visualized in the individual cortical map. Darker colors denote lower EV. The TPJ is shown as the region surrounded by the white line. The TPJ and connected regions within the DMN have low EV. Related to Figure 2 and Figure S1.

**Figure S4.**
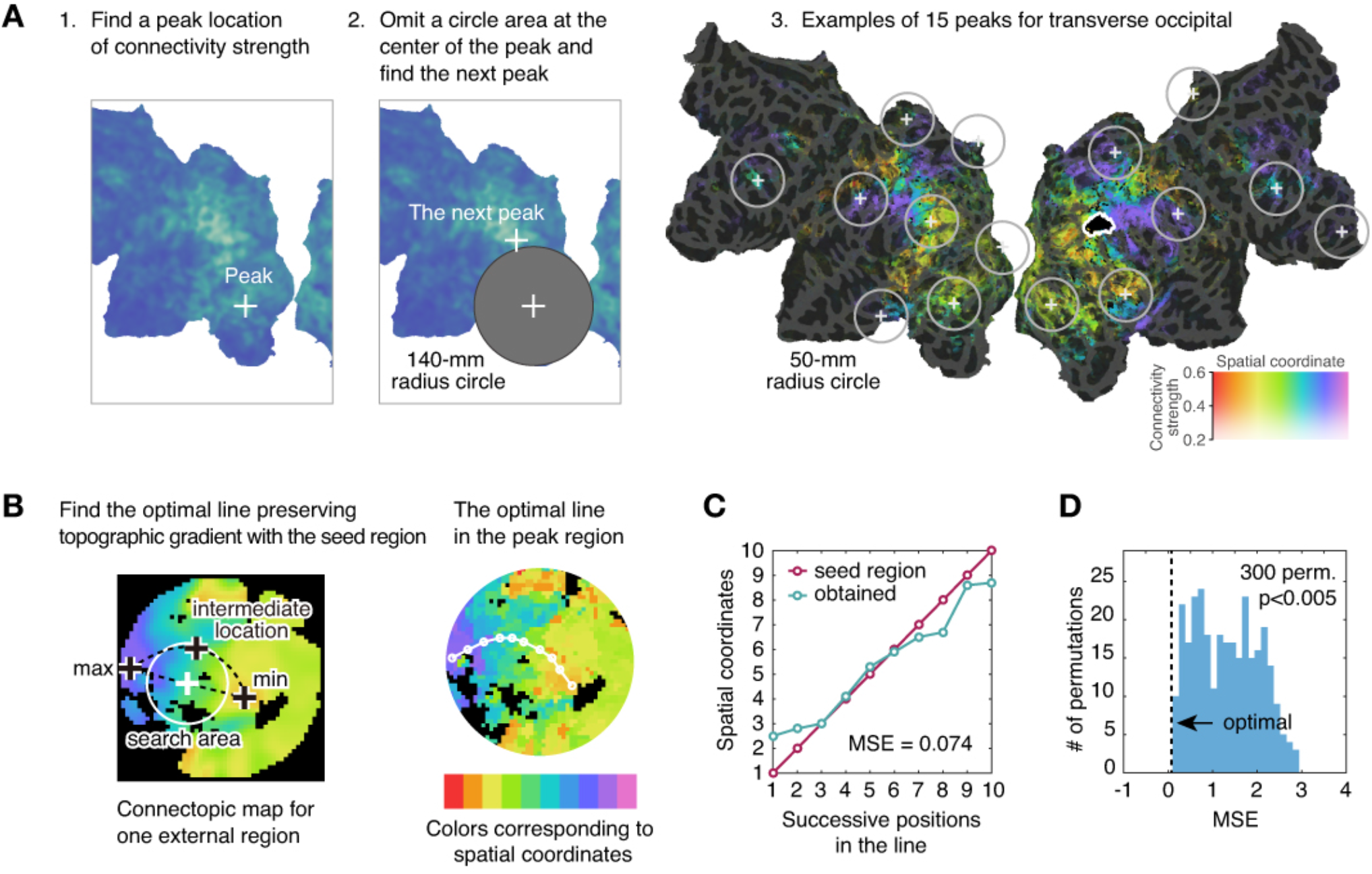
Procedure for calculating an MSE between the spatial coordinates for each of 156 anatomically-defined seed regions and those for each of the connected regions. (A) Workflow to identify 15 connected external regions (see Supplemental Information for details) (B) In each of the external regions, the optimal line was fitted to have a similar gradient to the seed topography from positions on the line. (C) Spatial coordinates obtained from the optimal line (turquoise) were compared with those in the seed region (red). Topographic dissimilarity between the two types of spatial coordinates was quantified using mean squared error (MSE). (D) Histogram of the MSEs for the randomly-defined lines (blue bins), and the MSE for the optimal line (black-dashed line). The MSE for the optimal line was significantly lower than the MSEs for the randomly-defined lines (*p* < 0.005, 300 permutations). Related to Figure 4.

**Figure S5.**
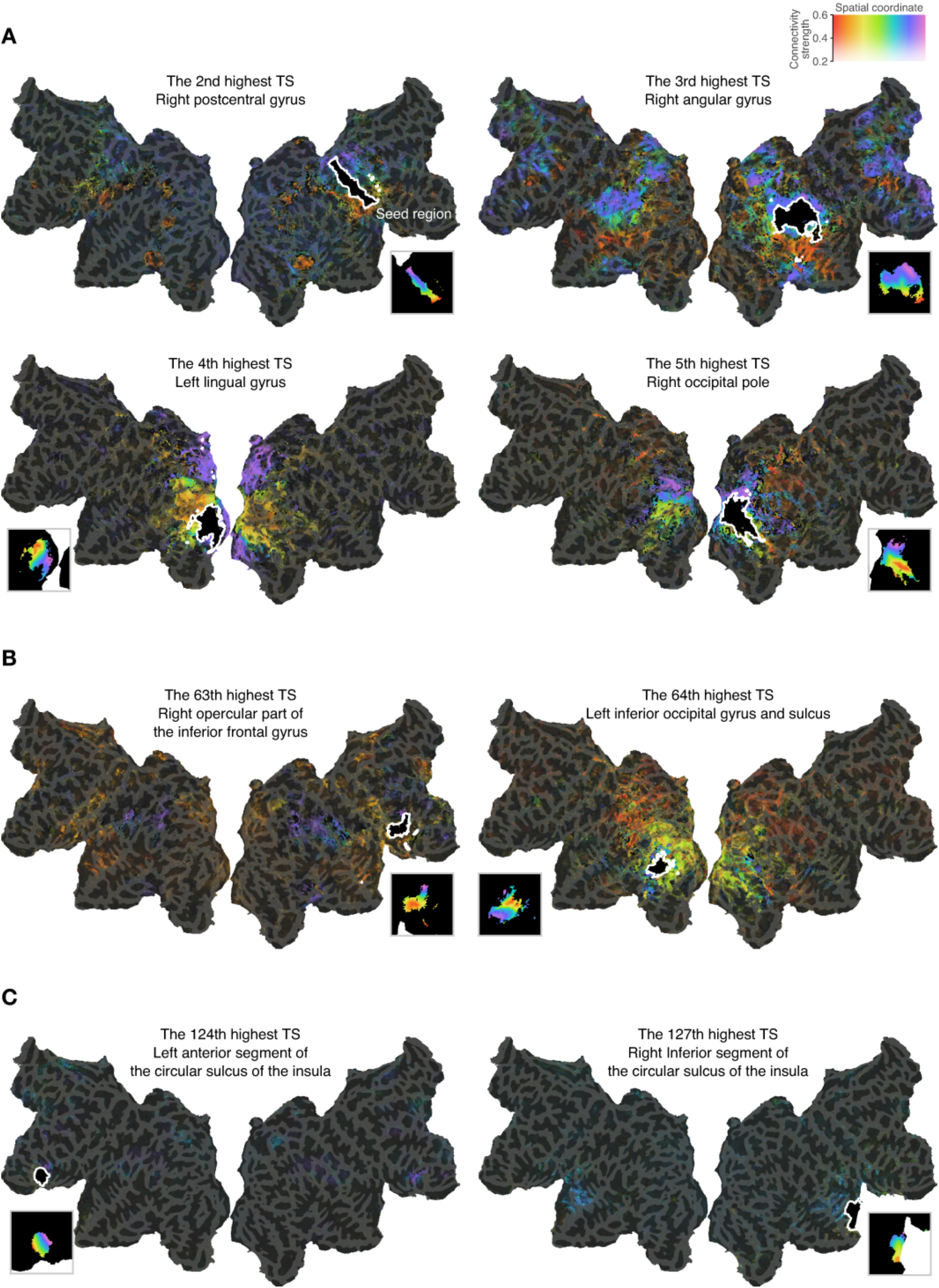
Examples of connectopic maps for a single subject S1 using seed regions showing high-, middle- and low-topographic similarity (TS) in connections with external regions. The seed regions were detected from the 156 anatomically-defined seed regions. (A) Connectopic maps for high-TS seed regions. In each map, we can observe similar topographic gradients in the seed region and the connected external region, while similar gradients were not clearly observed in (B) the middle-TS seed regions and not observed in (C) the low-TS seed regions. Related to Figure 4.

## Supplementary Methods

In our method, we automatically detected the fcTO-preserving seed regions from 156 labels as anatomically defined by the Destrieux atlas [Destrieux et al., 2010]. The abbreviation labels of the 156 regions are as follows (See Destrieux et al., 2010 to identify the label names corresponding to the abbreviation labels). ctx_lh_G_and_S_frontomargin, ctx_lh_G_and_S_occipital_inf, ctx_lh_G_and_S_paracentral, ctx_lh_G_and_S_subcentral, ctx_lh_G_and_S_transv_frontopol, ctx_lh_G_and_S_cingul-Ant, ctx_lh_G_and_S_cingul-Mid-Ant, ctx_lh_G_and_S_cingul-Mid-Post, ctx_lh_G_cingul-Post-dorsal, ctx_lh_G_cingul-Post-ventral, ctx_lh_G_cuneus, ctx_lh_G_front_inf-Opercular, ctx_lh_G_front_inf-Orbital, ctx_lh_G_front_inf-Triangul, ctx_lh_G_front_middle, ctx_lh_G_front_sup, ctx_lh_G_Ins_lg_and_S_cent_ins, ctx_lh_G_insular_short, ctx_lh_G_occipital_middle, ctx_lh_G_occipital_sup, ctx_lh_G_oc-temp_lat-fusifor, ctx_lh_G_oc-temp_med-Lingual, ctx_lh_G_oc-temp_med-Parahip, ctx_lh_G_orbital, ctx_lh_G_pariet_inf-Angular, ctx_lh_G_pariet_inf-Supramar, ctx_lh_G_parietal_sup, ctx_lh_G_postcentral, ctx_lh_G_precentral, ctx_lh_G_precuneus, ctx_lh_G_rectus, ctx_lh_G_subcallosal, ctx_lh_G_temp_sup-G_T_transv, ctx_lh_G_temp_sup-Lateral, ctx_lh_G_temp_sup-Plan_polar, ctx_lh_G_temp_sup-Plan_tempo, ctx_lh_G_temporal_inf, ctx_lh_G_temporal_middle, ctx_lh_Lat_Fis-ant-Horizont, ctx_lh_Lat_Fis-ant-Vertical, ctx_lh_Lat_Fis-post, ctx_lh_Pole_occipital, ctx_lh_Pole_temporal, ctx_lh_S_calcarine, ctx_lh_S_central, ctx_lh_S_cingul-Marginalis, ctx_lh_S_circular_insula_ant, ctx_lh_S_circular_insula_inf, ctx_lh_S_circular_insula_sup, ctx_lh_S_collat_transv_ant, ctx_lh_S_collat_transv_post, ctx_lh_S_front_inf, ctx_lh_S_front_middle, ctx_lh_S_front_sup, ctx_lh_S_interm_prim-Jensen, ctx_lh_S_intrapariet_and_P_trans, ctx_lh_S_oc_middle_and_Lunatus, ctx_lh_S_oc_sup_and_transversal, ctx_lh_S_occipital_ant, ctx_lh_S_oc-temp_lat, ctx_lh_S_oc-temp_med_and_Lingual, ctx_lh_S_orbital_lateral, ctx_lh_S_orbital_med-olfact, ctx_lh_S_orbital-H_Shaped, ctx_lh_S_parieto_occipital, ctx_lh_S_pericallosal, ctx_lh_S_postcentral, ctx_lh_S_precentral-inf-part, ctx_lh_S_precentral-sup-part, ctx_lh_S_suborbital, ctx_lh_S_subparietal, ctx_lh_S_temporal_inf, ctx_lh_S_temporal_sup, ctx_lh_S_temporal_transverse, ctx_rh_G_and_S_frontomargin, ctx_rh_G_and_S_occipital_inf, ctx_rh_G_and_S_paracentral, ctx_rh_G_and_S_subcentral, ctx_rh_G_and_S_transv_frontopol, ctx_rh_G_and_S_cingul-Ant, ctx_rh_G_and_S_cingul-Mid-Ant, ctx_rh_G_and_S_cingul-Mid-Post, ctx_rh_G_cingul-Post-dorsal, ctx_rh_G_cingul-Post-ventral, ctx_rh_G_cuneus, ctx_rh_G_front_inf-Opercular, ctx_rh_G_front_inf-Orbital, ctx_rh_G_front_inf-Triangul, ctx_rh_G_front_middle, ctx_rh_G_front_sup, ctx_rh_G_Ins_lg_and_S_cent_ins, ctx_rh_G_insular_short, ctx_rh_G_occipital_middle, ctx_rh_G_occipital_sup, ctx_rh_G_oc-temp_lat-fusifor, ctx_rh_G_oc-temp_med-Lingual, ctx_rh_G_oc-temp_med-Parahip, ctx_rh_G_orbital, ctx_rh_G_pariet_inf-Angular, ctx_rh_G_pariet_inf-Supramar, ctx_rh_G_parietal_sup, ctx_rh_G_postcentral, ctx_rh_G_precentral, ctx_rh_G_precuneus, ctx_rh_G_rectus, ctx_rh_G_subcallosal, ctx_rh_G_temp_sup-G_T_transv, ctx_rh_G_temp_sup-Lateral, ctx_rh_G_temp_sup-Plan_polar, ctx_rh_G_temp_sup-Plan_tempo, ctx_rh_G_temporal_inf, ctx_rh_G_temporal_middle, ctx_rh_Lat_Fis-ant-Horizont, ctx_rh_Lat_Fis-ant-Vertical, ctx_rh_Lat_Fis-post, ctx_rh_Pole_occipital, ctx_rh_Pole_temporal, ctx_rh_S_calcarine, ctx_rh_S_central, ctx_rh_S_cingul-Marginalis, ctx_rh_S_circular_insula_ant, ctx_rh_S_circular_insula_inf, ctx_rh_S_circular_insula_sup, ctx_rh_S_collat_transv_ant, ctx_rh_S_collat_transv_post, ctx_rh_S_front_inf, ctx_rh_S_front_middle, ctx_rh_S_front_sup, ctx_rh_S_interm_prim-Jensen, ctx_rh_S_intrapariet_and_P_trans, ctx_rh_S_oc_middle_and_Lunatus, ctx_rh_S_oc_sup_and_transversal, ctx_rh_S_occipital_ant, ctx_rh_S_oc-temp_lat, ctx_rh_S_oc-temp_med_and_Lingual, ctx_rh_S_orbital_lateral, ctx_rh_S_orbital_med-olfact, ctx_rh_S_orbital-H_Shaped, ctx_rh_S_parieto_occipital, ctx_rh_S_pericallosal, ctx_rh_S_postcentral, ctx_rh_S_precentral-inf-part, ctx_rh_S_precentral-sup-part, ctx_rh_S_suborbital, ctx_rh_S_subparietal, ctx_rh_S_temporal_inf, ctx_rh_S_temporal_sup, ctx_rh_S_temporal_transverse, Thalamus, Caudate, Putamen, Pallidum, Brain Stem, Hippocampus, Amygdala, Accumbens

